# Evolutionary effects of individual variation and dimensionality of higher-order interactions on robustness of species coexistence

**DOI:** 10.1101/2022.12.22.521465

**Authors:** Gaurav Baruah, György Barabás, Robert John

## Abstract

Although the eco-evolutionary effects of individual variation for species coexistence are still widely debated, theoretical evidence appears to support a negative impact on coexistence. Mechanistic models of eco-evolutionary effects of individual variation focus largely on pairwise interactions, while the dynamics of communities where both pairwise and higher-order interactions (HOIs) are pervasive are not known. In addition, most studies have focused on effects of high dimensional HOIs on species coexistence when in reality such HOIs could be highly structured and low-dimensional, as species interactions could primarily be mediated through phenotypic traits. Here, combining quantitative genetics and Lotka-Volterra equations, we explored the eco-evolutionary effects of individual variation on the patterns of species coexistence in a competitive community dictated by pairwise interactions and HOIs. Specifically, we compare six different models in which HOIs were modelled to be trait-mediated (low-dimensional) or random (high-dimensional) and evaluated its impact on robustness of species coexistence in the presence of different levels of individual variation. Across the six different models, we found that individual variation did not promote species coexistence, irrespective of whether interactions were pairwise or were of higher-order. However, individual trait variation could stabilize communities to external perturbation more so when interactions were of higher order. When compared across models, species coexistence is promoted when HOIs strengthen pairwise intraspecific competition more so than interspecific competition, and when HOIs act in a hierarchical manner. Additionally, across the models, we found that species’ traits tend to cluster together when individual variation in the community was low. We argue that, while individual variation can influence community patterns in many different ways, they more often lead to fewer species coexisting together.

## 1. Introduction

Explanations for multi-species coexistence in ecological communities have largely been sought at the species level by emphasizing mean differences among species driven by competitive interactions or life-history trade-offs (Violle & Jiang 2009; Clark 2010; Gravel *et al*. 2011; Kraft *et al*. 2015; Valladares *et al*. 2015). The idea being that differences among species along multiple ecological dimensions would minimize niche overlap and permit long-term species coexistence (Barabás & D’Andrea 2016). However, species appear to compete for only a small number of limiting resources (Hutchinson 1961; Laird & Schamp 2006; Shoresh *et al*. 2008; Li & Chesson 2016; Letten & Stouffer 2019) and numerous species may coexist despite little differences in demographic or resource-based niches (Condit *et al*. 2006), which poses a challenge for classical coexistence models.

Classical competition models of coexistence consider interactions among species pairs and require precise parameter trade-offs to stabilize communities or to limit the strength of competition among coexisting species in accordance with the competitive exclusion principle (Barabás *et al*. 2012). Theoretical studies with competition models further show that any stability achieved through pairwise competitive interactions can be disrupted by random interactions among species, and the number of coexisting species then declines inversely with the strength of their effective pairwise interactions (Bairey *et al*. 2016). Interaction strength, therefore, places an upper bound on the numbers of coexisting species, implying that strong pairwise competitive interactions alone cannot promote species coexistence in a large community with randomly interacting species pairs.

Interactions among species are not always constrained to species pairs and can involve higher-order combinations (Wilson 1992; Laird & Schamp 2006; Grilli *et al*. 2017; Mayfield & Stouffer 2017), where interactions between a species pair is modulated by one or more species (Fig. 1). In an ecological system where pairwise interactions structure communities, indirect or higher-order effects may modify these interactions and restructure communities (Levine *et al*. 2017). For example, a superior competitor for a limiting resource will inhibit an inferior competitor for the same resource, but a third species may modulate the strength of this inhibition without affecting either of the two competitors directly (Bairey *et al*. 2016). Such attenuation of the pairwise inhibitory effect can be density-mediated or trait-mediated, and can lead to qualitatively different community dynamics compared to the case with pure pairwise interactions. HOIs that have been studied thus far are mostly parameterized to be high-dimensional. In such high-dimensional HOIs, i.e., interaction coefficients can range from very low to high values and species coexistence could be promoted provided they fulfill certain conditions (Singh & Baruah 2021). By contrast, low-dimensional HOIs could come into effect when interactions are structured by just a handful of species traits. Indeed, in large communities, species interactions could be mediated by traits species possess (Guimarães *et al*. 2011; Maruyama *et al*. 2014; McPeek 2017) which consequently could structure communities and impact stability (Barabas & D’Andrea 2016; Baruah 2022). While the importance of such HOIs has been recognized (Levine *et al*. 2017), the nuances of such dimensionality of HOIs and networks of competitive pairwise interactions in communities have proven difficult to study theoretically or empirically.

**Figure 1:**
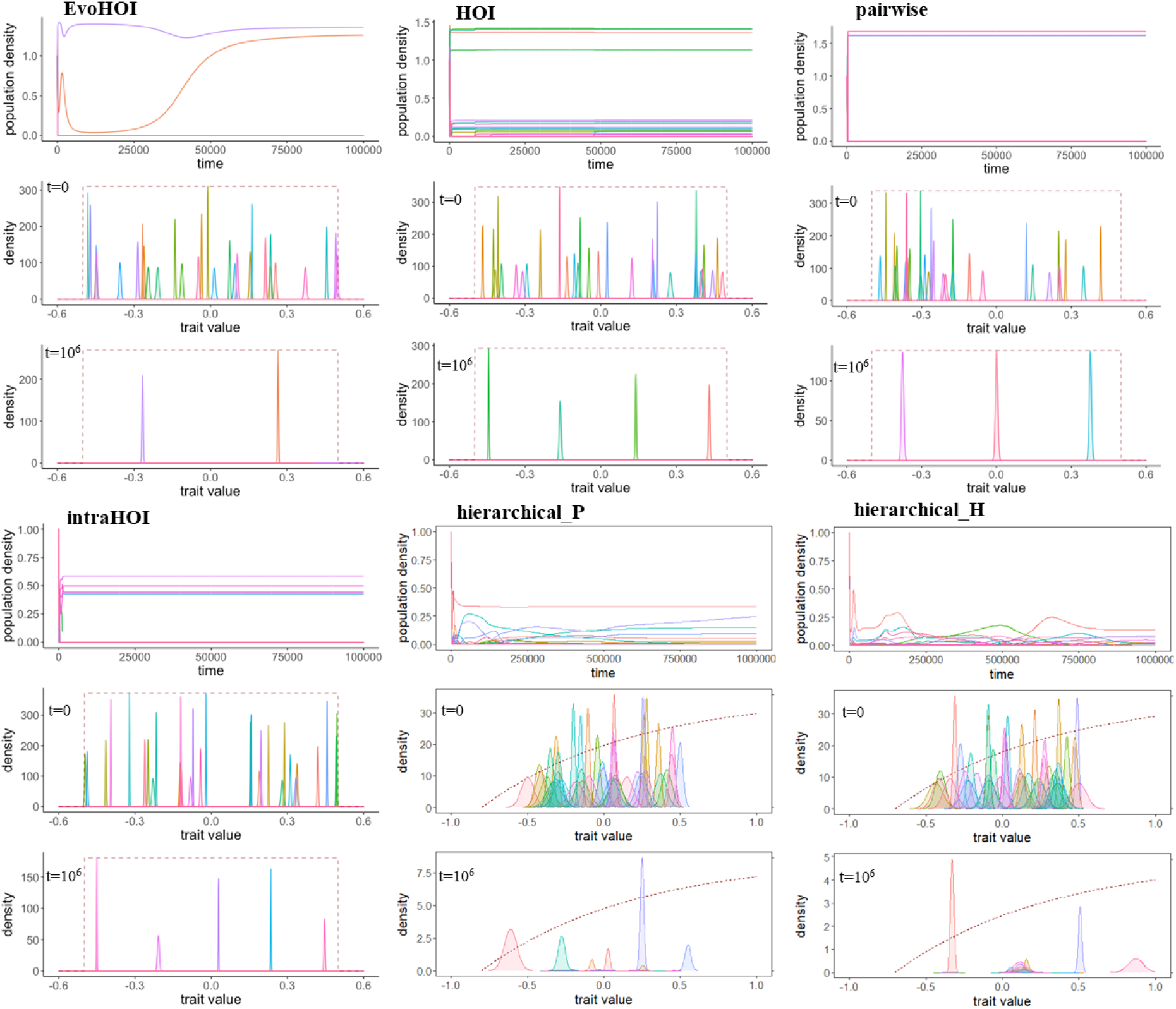
Eco-evolutionary dynamics of species competing in a trait axis with six different types of species interactions: pairwise, EvoHOI (evolutionary HOIs), HOI (mixed HOIs), intraHOIs (intraspecific HOIs stronger than interspecific HOIs), hierarchical pairwise model (hierarchical_P) and hierarchical higher-order model (hierarchical_H). Note that pairwise and evolutionary HOIs leads to lower number of species coexisting in comparison to when species interactions were dominated by stronger intraspecific HOIs or mixed HOIs. Here in the six models we show population dynamics as timeseries and snapshots of phenotypic trait distributions along the uni-dimensional axis for time points *t =* 0 and *t =* 10^6^. Red dashed line in subplots of trait distributions represent how rate of growth varies as trait value varies. Parameters used of the simulations: σ *=* [ 0.01, 0.05], ω *=* 0.1 for pairwise and EvoHOI models. For HOI, σ *=* [ 0.01, 0.05], ω *=* 0.1. HOIs were randomly sampled from U[-0.01,0.01]. For IntraHOIs, we ensured *∈*_*iik*_ to be strictly stronger in magnitude than interspecific HOIs *∈*_*ijk*_, *i* ≠ *j* ≠ *k*. This was done by sampling intraspecific HOIs from a random uniform distribution in the range [0.05, 0.1] and interspecific HOIs from a random uniform distribution in the range [0,0.04]. For hierarchical pairwise model, *=* [ 0.01, 0.05], ω *=* 0.1, and z_0_ *=* 0.7. similarly, for hierarchical higher order model, σ *=* [ 0.01, 0.05], ω *=* 0.1, and z_0_ *=* 0.7. Initial densities of all species was fixed at 0.5, initial trait values were uniformly sampled from the range [-0.5, 0.5] and heritability of the mean phenotypic trait was fixed for all species at 0.25.

A further consequence of emphasizing species-level mean differences to explain coexistence is that within-species or individual level variation has largely been ignored in species coexistence mechanisms (Siefert 2012; Hart *et al*. 2016). Empirical studies show that variation within species often exceeds the differences in species-level averages, but the consequences of intraspecific variation for species coexistence are yet to be fully explored (Barabás & D’Andrea 2016; Hausch *et al*. 2018). It is now clear that intraspecific variation can have both ecological and evolutionary effects on competitive interactions, which ultimately determine patterns of species coexistence (Pastore *et al*. 2021; Yamamichi *et al*. 2022). For example, intraspecific trait variation can undermine species coexistence by increasing or decreasing competitive ability, niche overlap, and even-spacing among species (Barabas & D’Andrea 2016), or by altering competitive outcomes through non-linear averaging of performances (Hart *et al*. 2016). There is equally compelling evidence that intraspecific variation promotes species coexistence, mainly by disrupting interspecific competitive abilities and attenuating the effects of strongly competitive individuals in a community (Bolnick *et al*. 2011). Experimental work has shown that intraspecific variation although allows a community to be resilient to invaders, creates the opportunity for competitive exclusion among strong competitors (Hausch *et al*. 2018). Empirical studies in the field show, for instance, that leaf economic traits such as specific leaf area, leaf N and P consistently display high intraspecific variation (Meziane & Shipley 1999; Vasseur *et al*. 2012), and influence community assembly and ecosystem functioning (Reich 2014). Given that high levels of intraspecific trait variation within communities appears to be more a rule than an exception, the implications of intraspecific variation to community structure merits detailed investigation.

It is clear that both higher-order species interactions and intraspecific variation can have significant influence on community structure, but these two processes have always been investigated separately in studies on species coexistence. The effects of intraspecific trait variation and eco-evolutionary dynamics on structuring large communities where both pairwise and HOIs dominate a community are therefore unknown. Theoretical analyses indicate that purely pairwise interactions in a community lead to even trait spacing when intraspecific variation is high. Consequently, high intraspecific variation should promote competitive exclusion of inferior species in a large community. However, a community dominated by both pairwise and HOIs could show reduced even spacing of species in a trait axis and greater trait clustering. The apparent mechanism could be that with intraspecific variation present, HOIs could significantly alleviate and stabilize the negative pairwise interactions that would otherwise lead to distinct spacing. It therefore appears valuable to link HOIs and intraspecific variation in order to understand their combined influence on species coexistence and community structure.

In this study, we examine the importance of dimensionality of HOIs and intraspecific variation in structuring species coexistence and trait patterning. We do this by reconciling quantitative traits with a Lotka-Volterra modelling approach, where the dynamics of the whole community are mediated both by pairwise competitive interactions as well as higher-order three-way interactions. Specifically, we model a one-dimensional quantitative trait that contributes to the pairwise competitive ability of species interacting in the community. Next, we formulate six models of how distinct dimensions of HOIs could influence pairwise competition. While modelling phenotypic-mediated species competition we do not restrict to Gaussian based interaction kernels but also consider kernels that could be hierarchical in nature. On that note, species with specific trait value could have a competitive advantage over others just because of their position in the trait axis that simultaneously trades-off with growth rate similar to the model of Adler & Mosquera 2000. Overall, we found that the impact of individual trait variation on robustness of species coexistence was dependent on the dimensionality of species interactions and how species compete with each other. Our study links recent ecological studies of HOIs with eco-evolutionary dynamics and intraspecific variation.

## 2. Methods and Models

### 2.1 Model 1: Multispecies model with just pairwise interactions

In our community model, we consider species competing with each other along a one-dimensional trait axis. Two phenotypes with trait values *z* and *z’* along the axis compete with one another depending on the distance of their traits: more similar phenotypes compete more strongly. Species are composed of phenotypes which, in line with quantitative genetic principles (Barton *et al*. 2017), are normally distributed with mean *u*_*i*_ and variance 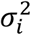 for species *i*. Under such conditions, the dynamics of species densities are given by Lotka-Volterra equations as (Barabas & D’Andrea 2016):

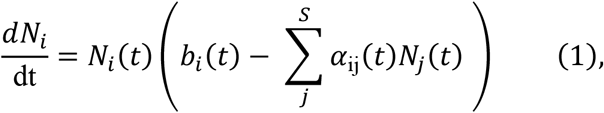

And the dynamics of the mean competitive trait *u*_*i*_ is given by:

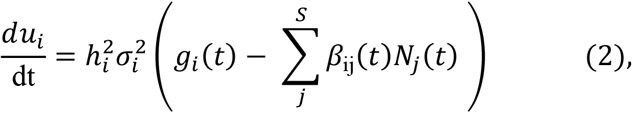

where *α*_ij_(t) describes the pairwise competition coefficient of species *i* with species *j* at any time *t*. This competition coefficient derives directly from a Gaussian competition kernel (Appendix 2). If the two species are similar to each other in terms of their average trait value *u*, then competition between them is stronger than when they are farther apart in the trait axis; 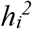 is the heritability of the mean trait for species *i, b*_*i*_(t) describes the growth rate of the species *i* in the absence of any competition which is determined by where they lie in the trait axis *z*; *g*_*i*_(t) describes selection on the trait stemming from the differential growth of the phenotypes comprising the species, and *β*_ij_(t) quantifies the evolutionary pressure on the trait *z* for species *i* due to competition with the species *j* in the community (this has been derived in Barabás and D’Andrea 2016).

### 2.2 Model 2: Evolutionary trait-based higher-order interactions (EvoHOI)

Equations 1 and 2 capture the eco-evolutionary dynamics of a multispecies community where pairwise interactions dominate community dynamics. It is plausible that such a community exhibits higher-order interactions as well, rather than just pairwise ones. In extension to the above model, we include trait-mediated three-way higher-order interactions where the traits of three species influence pairwise competitive interactions. HOIs and evolutionary dynamics of species coexistence have not been considered together, until now. Under these circumstances, the equations become (Appendix 2):

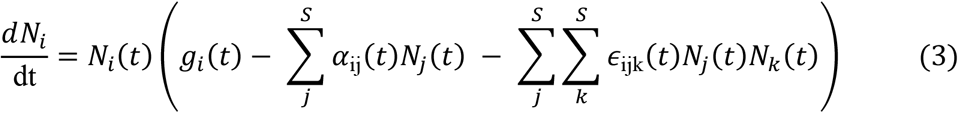

And the dynamics of the mean trait *u*_*i*_ are given by:

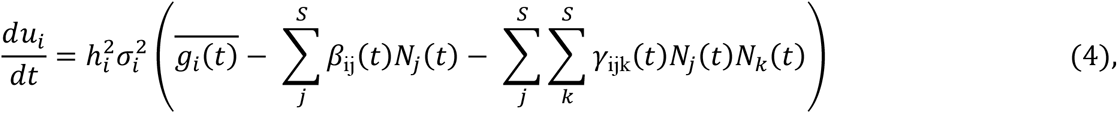

where *∈*_ijk_ gives the 3-way interactions that are density mediated HOIs (*∈*_ijk_(t) is termed inter-specific HOIs, and *∈*_iik_(t) and *∈*_iii_(t) are termed as intraspecific HOIs) (Mayfield & Stouffer 2017; Letten & Stouffer 2019); *γ*_ijk_ denotes 3-way interactions (*γ*_ijk_ as interspecific evolutionary effects; *γ*_iii_ and *γ*_iij_ as intraspecific evolutionary HOI effects)(Singh & Baruah 2021; Kleinhesselink *et al*. 2022) affecting the evolutionary dynamics of mean trait *u*_*i*_ for species *i*. Similar to the pairwise Gaussian interaction kernel, the three-way interaction remains Gaussian with a third species *k* influencing the interaction between the two species *i* and *j* given as 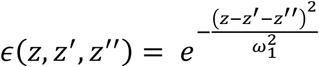. *∈*_ijk_(t) and γ_ijk_(t) are three-index tensors of size (*S* × *S* × *S*), where *S* is the total number of species in the community derived after integrating the three-way Gaussian kernel with respect to the trait distributions of three species interacting (Appendix 2). 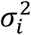 and 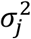 are the intraspecific trait variation for species *i* and species *j* respectively; 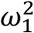 is the width of the competition kernel which was Gaussian (Appendix 2); *u*_*i*_(t) is the average trait value of species *i* and *u*_*j*_(t) is the average trait value for species *j*. Thereby, eco-evolutionary dynamics in this purely competitive community is dominated not only by pairwise trait-based competition but also by three-way higher-order interactions. In such a case, eco-evolutionary dynamics might deviate from dynamics dominated by purely pairwise competitive coefficients as in Barabás and D’Andrea (2016). For details of the formulation see Appendices 1-2. To be noted that higher-order effects on competitive ability of species are mediated primarily through the one-dimensional mean phenotypic trait. As such, such HOIs are (EvoHOI) are low dimensional in nature, as their effects are bounded by the trait position of a species as well as the by the variance of the trait.

### 2.3 Model 3: Evolutionary hierarchical trait-based HOI (Heirarchical)

We model another trait-mediated HOI where interactions among species were also mediated through mean phenotypic traits. However, the pairwise and higher-order interaction kernel was not Gaussian but could be written as, 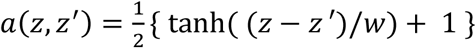 and 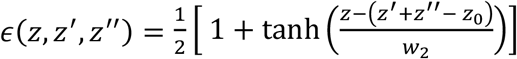 respectively. Here, *a*(*z, z*′) and *∈*(*z, z*′, *z*′′) represent pairwise competition and higher-order competition kernel. *z, z*′ are phenotypic trait values of two individuals engaged in pairwise competitive kernel in *a*(*z, z*′), and *z, z*’, *z*’’ represent phenotypic trait values of three individuals engaged in three-way interaction in *ε*(*z, z*′, *z*′′). *w* and *w*_2_ controls the strength of pairwise and three-way higher-order interaction. The pairwise competitive kernel indicates that competition is hierarchical (Adler & Mosquera 2000). For instance, larger body sized individuals could dominate smaller individuals in acquiring resources. This however comes with a trade-off with rate of growth such that competitively superior individuals trades off with low growth rate. The HOI interaction kernel written above was defined such that if two individuals compete with each other a third individual can modulate this pairwise competition in a way that could benefit the weaker competitor. Biologically this translates to a scenario for instance when a superior competitor dominates an inferior competitor for resources, but an another competitor could modulate this pairwise interaction in a way that benefits the weaker individual (van Veen *et al*. 2005; Abrams 2007). In this particular model, there is a trade-off with being a better competitor and having a higher growth rate. Species with higher phenotypic trait value which confers them a higher competitive ability trades off with lower intrinsic average growth rate given as:

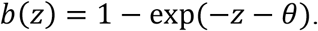

Thus governing population dynamics and trait dynamics can be similarly written as equation 3 and 4 (see details regarding the derivation in the appendix 1). With this model, we specifically evaluate how individual variation affects species coexistence in only pairwise model devoid of any HOI and contrast it with pairwise and HOI. In the case of pure pairwise interactions, *∈*(*z, z*′, *z*′′) = 0 and dynamics is dictated only by competitive pairwise *a*(*z, z*′). Note that since HOI interactions were constrained due to phenotypes of species in the uni-dimensional trait axis, such HOIs are low dimensional in nature regardless of the hierarchical nature of competitive interactions.

### 2.4 Model 3: Intraspecific higher-order interactions (intraHOI)

We should state here that three-way HOIs need not be trait-mediated, and that HOIs could also be represented by, for example, interactions that were dependent on the environment or other traits. The model 2 represented by trait-mediated HOIs were low-dimensional and structured by how species compete for resources along a one-dimensional trait axis. On the other hand, when HOIs were modelled to be independent of such phenotypic traits, and represented by random coefficients such that their effects have a wider distribution, they become high-dimensional in nature. However, there is little empirical understanding of such interactions in nature, and here we chose random HOIs that directly influence species densities but have no direct impact on the evolutionary dynamics of the mean species trait values. In this particular case, HOIs will only indirectly impact the evolution of mean species traits through changes in species densities. This means that if individual variation did influence patterns of species coexistence with this particular formulation of HOIs, then it would be the result of the combined effect of both variation in mean traits and high-dimensional HOIs. Specifically, we wanted to evaluate how the structure of HOIs could influence robustness of species coexistence and trait patterning. Therefore, in this specific model, when sampling *∈*_ijk_ we ensured intraspecific HOIs *∈*_*iik*_ to be strictly stronger in magnitude than interspecific HOIs *∈*_*ijk*_, *i* ≠ *j* ≠ *k*. This was done by sampling intraspecific HOIs from a uniform distribution in the range [0.05, 0.1] and interspecific HOIs from a uniform distribution in the range [0, 0.04], with 40 independent replicates. This formulation ensured that HOIs strengthened pairwise intraspecific competition more so than they strengthened interspecific competition (Singh & Baruah 2021).

### 2.3 Model 4: Mixed higher-order interactions (HOI)

In this final model, we ensured that there was no specific structure to HOIs. Specifically, we randomly sampled HOIs, *∈*_*ijk*_, from a uniform distribution within the range [-0.01, 0.01] with 40 replicates. This meant that HOIs impacting species densities could both intensify as well as alleviate competitive interactions, since HOIs also could take positive values and thereby alleviate pairwise competitive interactions. Such HOIs are again high-dimensional in nature as their effects have a wider distribution and are not bounded by species mean traits or species trait variances in general.

### 2.4 Simulations of the model with higher-order interactions

We assessed the effect of different levels of intraspecific trait variation on community structure and species coexistence using data generated from simulations of our community models. We simulated both trait dynamics and population dynamics resulting from equations (3) and (4). We started each simulation with forty species and all the forty species were randomly assigned an initial mean trait value in the range −0.5 to 0.5 on the one-dimensional trait axis. Outside this trait regime, the fitness value of a species is set to be extreme, with negative growth rate, as given by the growth kernel (Appendix 1). This, however, only applied to all the models except model 3 i.e., the hierarchical model. Biologically, this strict criterion means that outside this trait boundary resource acquisition by a species would be too low to have a positive growth rate. For the hierarchical model, growth rate increases as trait value increases. We carried out 40 replicate simulations for each level of intraspecific variation. We also simultaneously tested the influence the strength of pairwise competition denoted by the width of the Gaussian pairwise competition kernel *ω* and slope *ω* for the hierarchical pairwise *tanh*() function using a fully factorial design where all possible combinations of intraspecific variation and interaction strength were tested for their influence on species coexistence. In all our simulations, heritability 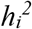of the trait for all species was fixed at 0.25 and we didn’t vary the strength of trait-based HOIs or hierarchical HOIs given by *ω*_1_ and *ω*_2_respectively which was fixed at 0.1. We evolved our community till it reached equilibrium. To be noted that species richness could be sensitive to extinction thresholds, hence we also measured species diversity as inverse Simpson’s diversity index. At the end of each replicate simulation, we measure species diversity as inverse Simpson’s diversity index given as 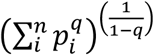, where *q* = 2, *p*_*i*_ is the proportional abundance of species *i*. This diversity index takes into account the effective number of equally-abundant species.

### 2.5 Levels of width of the competition kernel and intraspecific variation

The width of the pairwise competition kernel *ω* was varied from 0.1, 0.2, and 0.3. For each *ω*, three different levels of intraspecific variation were tested in a fully factorial design (3 different *ω* values × 3 different σ^2^ values × 4 different models × 40 replicates). Specifically, for each *ω*, intraspecific variation for each of the 40 species in the community was randomly sampled from a uniform distribution with three different levels: a) low variation: σ *=* [ 0.01, 0.05]; b) intermediate variation: σ *=* [ 0.05, 0.1]; and c) high variation: σ *=* [ 0.1, 0.5] (See Table 1 for the parameters used).

**Table 1:** List of parameters values used in the model. Note that variables are not given any value, but parameters for which the model simulations are tested are given values

| Parameters | Description | Value |
| --- | --- | --- |
| $N_i$ | density of species $i$ | variable |
| $\theta$ | | 0.5 |
| $b_i$ | growth rate of species $i$; | $\frac{1}{2} [\text{erf}(\frac{\theta - \mu_i}{\sqrt{2}\sigma_i}) + \text{erf}(\frac{\theta + \mu_i}{\sqrt{2}\sigma_i})]$ |
| $\alpha_{ij}$ | competition coefficient between<br>species $i$ and species $j$ | $\frac{\omega}{\sqrt{2\sigma_i^2 + 2\sigma_j^2 + \omega^2}} \exp(\frac{-(\mu_i(t) - \mu_j(t))^2}{2\sigma_i^2 + 2\sigma_j^2 + \omega^2})$ |
| $\epsilon_{ijk}$ | Three-way competitive interaction:<br>species $i,j$ influenced by species $k$ | as given in equation |
| $u_i$ | mean trait value of species $i$ | values in range from -0.5 to 0.5,<br>Initial trait values are randomly assigned |
| $\overline{b_i}$ | Environmental cue at time $t$ | $\frac{1}{\sqrt{2\pi}\sigma_i} [\exp(\frac{-(\theta + \mu_i)^2}{2\sigma_i^2}) - \exp(\frac{-(\theta - \mu_i)^2}{2\sigma_i^2})]$ |
| $\gamma_{ijk}$ | Evolutionary pressure on trait value<br>of species $i$ | |
|  | due to three-way competitive interactions, such that pairwise competition between two species $i$ and $j$ is modulated by the third species $k$, (a tensor of size $S \times S \times S$) | as given in equation (see supplementary equation 7) |
| $\sigma_i, \sigma_j$ | Trait standard deviations of species $i, j$; trait variation $a$<br>re uniformly sampled for each replicate from theses ranges | a) High variation = [0.1,0.5]<br>b) medium variation = [0.05,0.1]<br>c) low variation = [0.001,0.05] |
| $S$ | Total number of species in the community; | 40 |
| $\omega$ | width of the competition kernel signifying strength in pairwise competition | 0.1, 0.2, 0.3 |
| $\omega_2, z_0$ | Strength of trait-based HOI interaction, upper bound of trait for hierarchical model respectively. | 0.1, 0.7 |

### 2.6 Trait clustering

Theoretical models have suggested that species coexisting together tend to spread more evenly along a trait axis than expected from random allocation (Barabas & D’Andrea 2016; D’Andrea & Ostling 2016), though evidence for this is mixed given the convoluted nature of the problem (D’Andrea & Ostling 2016). Here, we use a quantitative metric to evaluate the effect of intraspecific variation on the patterning of traits in the trait-axis. We measured trait similarity among coexisting species by measuring the coefficient of variation (CV) of adjacent trait means (D’Andrea & Ostling 2016). High values of CV would indicate clustering of trait means of species in the trait axis while lower CV values would indicate even spacing of traits. In addition, we also compared trait-clustering results for the five different models alongside three different levels of individual trait variation. To evaluate the significance of trait clustering we also used a null model consisting of trait values of species drawn from a uniform distribution matching the final species richness of communities from the six models. Next, we compared the CVs of the communities originating from the six models with the 1000 corresponding random communities and calculate the P-value as the proportion of null CVs lower than the CVs of the communities from the corresponding six models. Lower P-values would indicate that communities from the six models were more spaced in the trait axis than by chance.

### 2.7 Robustness of species coexistence

The (ecological) robustness of our community model with higher-order interactions was measured by calculating the Jacobian matrix of the ecological part of the dynamics at equilibrium (Appendix 4):

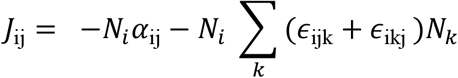

At the end of our simulations, a subset of the species in the original communities always stably coexisted, but some of these final communities were more robust against environmental perturbations than others. We measured the robustness of the community by taking the geometric mean of the absolute values of the eigenvalues of the Jacobian (Barabás & D’Andrea 2016; Appendix 4). For *S* species, this is the same as the *S*th root of the absolute value of the matrix’s determinant. This quantity measures the (geometric) average return times in response to environmental perturbations. For each replicate simulation of each level of intraspecific variation, we calculated the average community robustness as the measure to evaluate how intraspecific variation affects the robustness of species coexistence.

## 3. Results

### 3.1 Effect of intraspecific variation and width of competition on species coexistence

We found that when interactions are purely pairwise, increase in intraspecific variation led to a decrease in species diversity. When evolutionary HOIs come into play, diversity further decreases as individual variation increases across all ω (Fig 2). In addition, when high-dimensional mixed HOIs (HOI) comes into play, species diversity was the highest when trait variation was lowest across three different levels ω (Fig. 2, model HOI). Similarly, for the intraHOI model, which is also high-dimensional, where HOIs strengthened pairwise intraspecific competition more than interspecific competition, species coexistence was highest when trait variation was lowest the lowest (Fig. 2). In similar manner, in the hierarchical pairwise (hierarchical_P) and hierarchical higher-order model (hierarchical_H), medium to low amount of trait variation led to high species diversity in comparison to scenarios of high trait variation.

**Figure 2:**
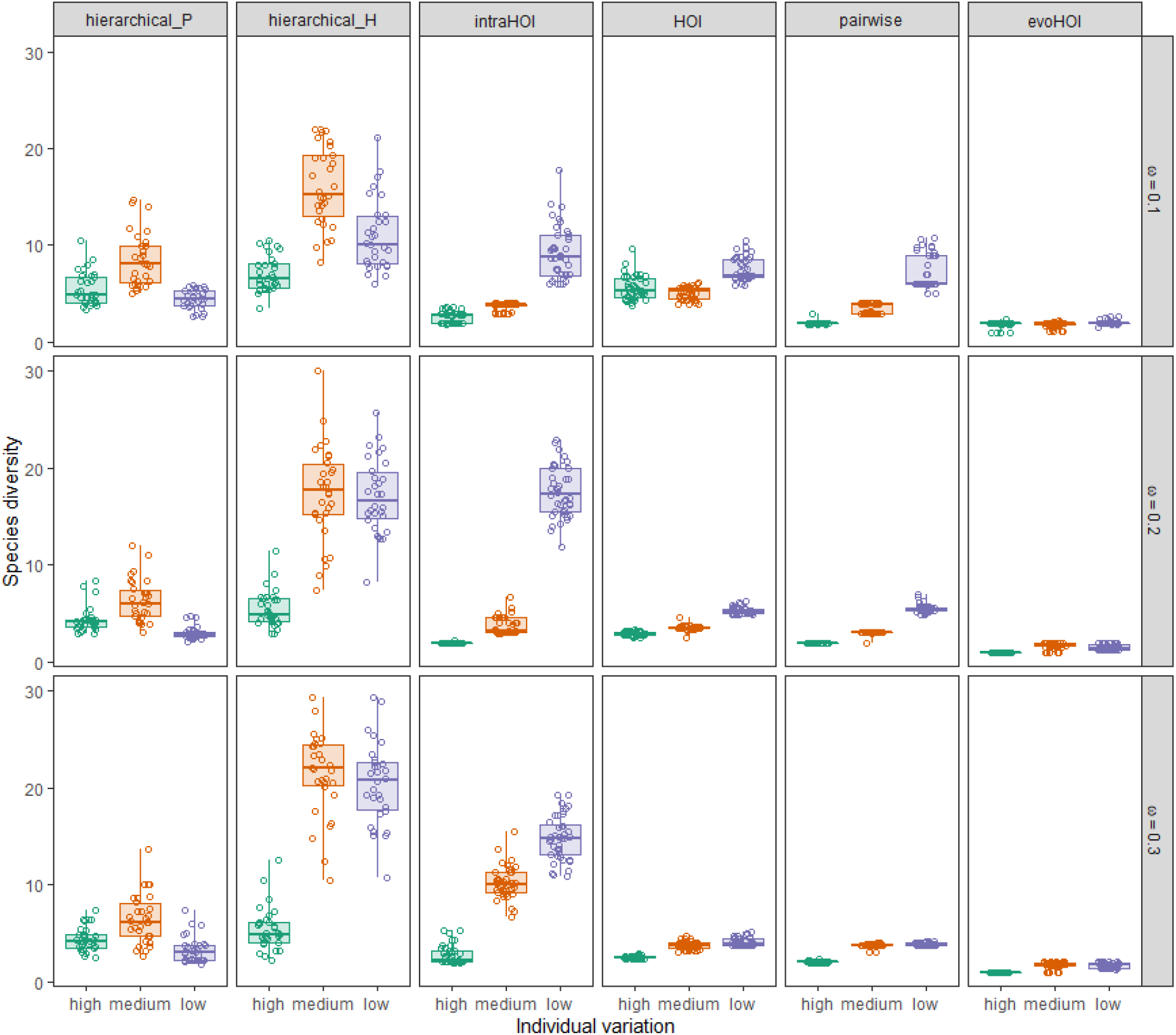
Effect of individual trait variation on species diversity measured as the inverse of Simpson’s diversity index. Boxplots denote the effective number of species that coexisted at the end of the simulations for different competition widths denoted by *ω* levels and for six different interaction models.

When comparing across different models, evolutionary HOIs, which were low-dimensional (evoHOI), disrupted species coexistence the most within every level of individual variation and across ω. However, when high-dimensional HOIs strengthened intraspecific competition (intraHOI), species coexistence was stabilized (across all *ω*), and when compared with other types of model. As competition width increased, the effect of intraHOI in conjunction with low individual variation on species coexistence increased (Fig. 2). Across all the models, low-dimensional trait-based HOI could still lead to high species coexistence provided higher-order interactions act in a hierarchical manner. Across all models, hierarchical trait-based HOI model lead to highest number of species coexisting in comparison to all other models.

### 3.2 Trait clustering

There was not much differences in trait clustering quantified by CV values across different levels of individual variation. Higher intraspecific variation led to slightly higher species clustering in EvoHOI and hierarchical higher-order models (hierarchical_H) consistently across different ω. In contrast, in case of intraHOI model, lower trait variation overall led to higher trait clustering especially at higher competitive width. For pairwise model, lower trait variation led to more species clustering as competitive width ω increased (Fig 3A).

**Figure 3:**
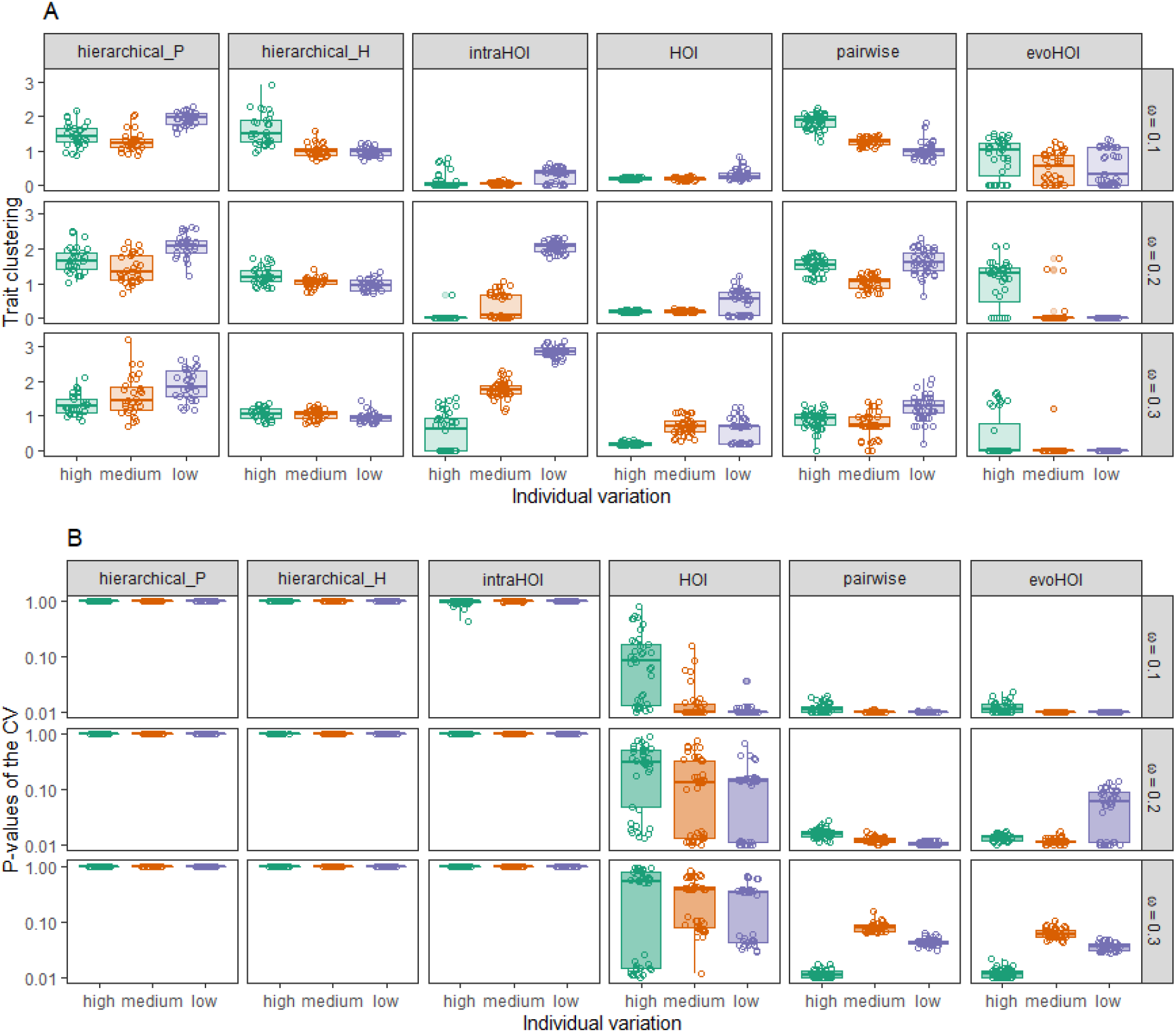
A) Effect of intraspecific variation on trait clustering (measured as coefficient of variation, CV, in final mean trait values of species that coexisted) for different levels of the competition width, *ω* and for the six different models. Higher values of trait clustering indicate higher trait overlap. Note that mean trait clustering increases with decreasing levels of intraspecific variation when intraspecific HOIs were stronger than interspecific HOIs (intraHOI). In the case of lower dimensional HOIs (evoHOI), trait clustering decreases as trait variation decreases. For the hierarchical models, high trait variation caused higher trait clustering for the hierarchical higher-order model in contrast to the pairwise hierarchical model. B) P-values of trait clustering indicating the probability that even spacing of species in the uni-dimensional trait axis is purely due to chance. Low p-values indicate unique species trait pattering. Lower dimensional HOI i.e., evoHOI and pairwise competition led to unique trait clustering. In contrast, hierarchical trait-based models whether it was pairwise (Hierarchical_P) or HOI (Hierarchical_H) led to trait patterning that had high p-values indicating that the degree of spacing could be produced by chance alone.

Trait patterning of species in the uni-dimensional axis was unique for the pairwise model and EvoHOI models across all the levels of trait variation and ω. Higher ω in these two models increases P-values (Fig 3B). This is due to the fact that higher ω increases competition which results in overall lower species diversity in comparison to lower ω which then reduces the statistical power while increasing the overall P-values. Surprisingly, for the higher-dimensional HOI models such as intraHOI and HOI, p-values were substantially larger indicating the communities produced might not be significantly different from random communities. One other surprising result was that, hierarchical models produced the highest p-values across all levels of trait variation and ω (Fig 3B).

### 3.3 Robustness of species coexistence

High-dimensional HOIs i.e., intraHOI and HOI models, exhibited higher average robustness in comparison to other models. Low-dimensional trait-based HOIs had lower average robustness across different competitive widths and trait variation in comparison to high-dimensional HOI models.

With increases in intraspecific variation across pairwise competitive width *ω*, average robustness of community coexistence increased, particularly for high-dimensional HOI models such as intraHOI and HOIs. Community robustness was highest when species exhibited high intraspecific variation and when interactions were HOIs but structured in a way that intraspecific HOIs were stronger than interspecific HOIs (Fig. 4, intraHOI). However, higher number of species coexisting led to a low community robustness, irrespecitive of whether species exhibited high or low variation. In specific models like IntraHOI, low trait variation led to higher number of species coexisting but at the cost of low community robustness. In contrast, high variation led to lower number of species coexisting but with high community robustness (Fig 4). Similarly, in models of pairwise interaction, HOI, evoHOI, higher trait variation could lead to slightly higher community robustness if such high trait variation led to less species coexisting at evolutionary equilibrium. In contrast, hierarchical pairwise and HOI model led to low community robustness across different levels of trait variation.

**Figure 4:**
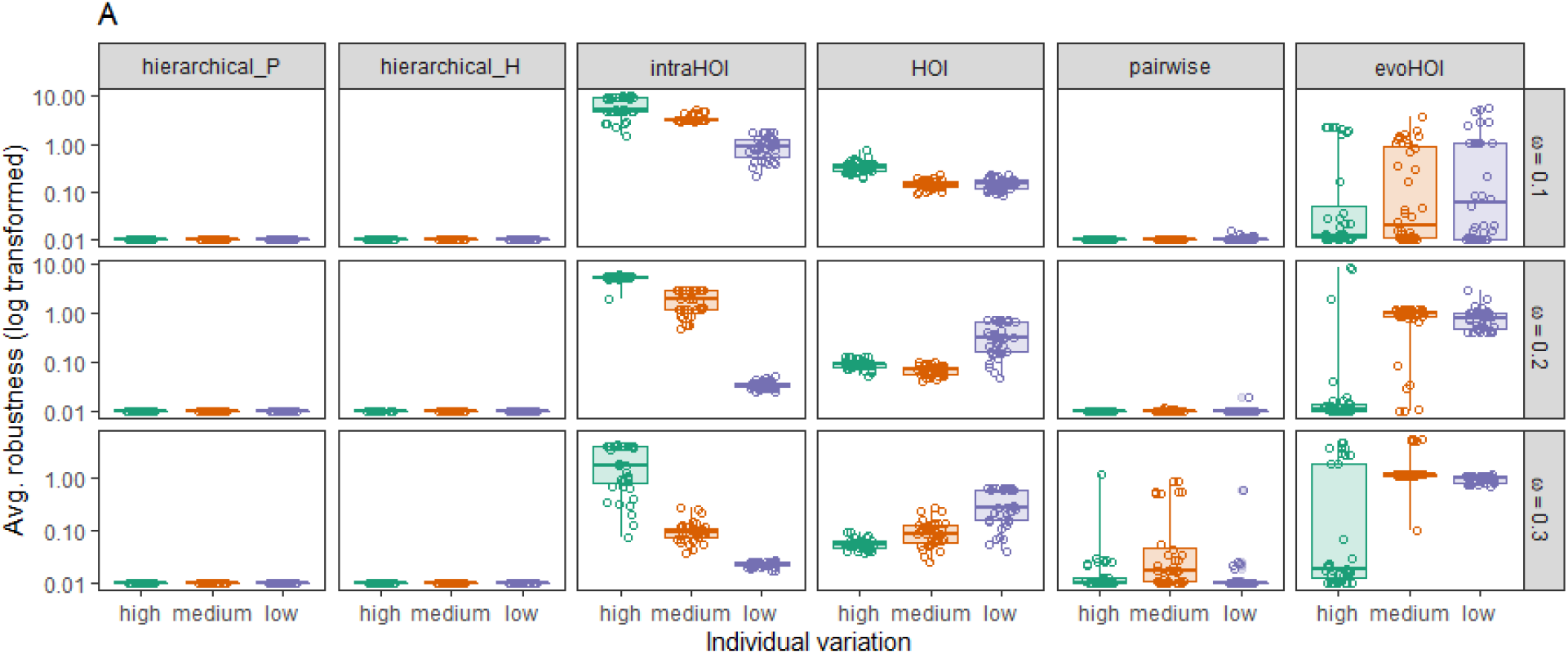
Effect of intraspecific variation on average community robustness for different levels of the competition width, *ω* and for six different interaction models.

## 4. Discussion

In mechanistic models of pairwise competition where interaction strengths vary with the density of both heterospecifics and conspecifics, HOIs emerge naturally as a suite of non-additive effects that influence competition between the focal species pair (Abrams 1983; Letten & Stouffer 2019). Where HOIs have been explicitly modelled in phenomenological ecological models, they could act as a stabilizing factor in maintaining species diversity (Bairey *et al*. 2016; Grilli *et al*. 2017). We modelled the evolution of a trait that dictates competitive ability between species and introduced higher-order competitive interactions where pairwise interactions were modulated by a third species. One of the key objectives of our study was to investigate the influence of intraspecific variation in trait values on species coexistence. We found that when intraspecific variation increased, species richness decreased, in a manner analogous to what Barabás and D’Andrea (2016) found (Fig.1, Fig. 2). In addition to this, dimensionality of higher-order and individual trait variation consequently determined the patterning of species coexistence on a one-dimensional trait axis. Furthermore, trait-based higher-order interactions could promote species coexistence provided the phenotypic trait of the third species impacts pairwise interaction in a hierarchical manner.

A strong competitor on the trait axis can negatively affect the growth of inferior competitors. Our results suggest that with the introduction of three-way interactions, this dominance of the competitively superior species and its impact on coexistence is dependent on the structure, dimensionality, and the manner in which HOIs impact pairwise competition. Specifically, in pairwise interacting species, when a third species strengthens intraspecific competition more than interspecific competition (intraHOIs), species coexistence was promoted, especially at higher levels of competition width. This was because the third species further restricts the growth of the other two species by increasing intraspecific pairwise competition (Singh & Baruah 2021). This is analogous to the pairwise coexistence rule where species must limit themselves more than limiting others in order for coexistence to prevail (Fig. 2). Such HOIs could be considered high-dimensional, as besides low-dimensional species competition based on species mean trait values, these HOIs are random coefficients sampled from a wider distribution than species mean trait values. Such higher-order effects could be a manifestation of variety of underlying unmeasured traits of species, or could be a result of variation in environmental conditions. Sampling specific random HOIs thus could fairly mimic such a scenario. Our results indicated that when such HOIs were not dictated by low-dimensional trait values but dictated by certain set of rules that impact pairwise competition, it could result in higher number of species coexisting. Trait-based HOIs, which are low-dimensional HOIs, were bounded by mean phenotypic trait values of species and trait variation. Trait-based HOIs acting in a Gaussian three-way competition kernel in a way i.e., evoHOI, further disrupts species coexistence. For instance, two species with similar phenotypic trait values will compete very strongly. In that case, if the third species also has a similar phenotypic trait value, it will further intensify pairwise competition. Consequently, this further disrupts species coexistence in contrast to when species only compete through pairwise manner. However, trait-based low dimensional HOIs could promote species coexistence. This could be possible if the three-way HOIs act in a hierarchical manner. In our hierarchical higher-order model, it was possible for a third species to alleviate competitive dominance of one species on another in a way that promoted species coexistence. In the hierarchical model, species with lower trait values are more competitively dominant than those with higher trait values. This however came with a trade-off – competitively dominant species had lower growth rate. With this framework, in the hierarchical pairwise model, high trait variation still led to low number of species coexisting. In contrast, in hierarchical trait-based HOI model, more species coexisted across all levels of strength of pairwise competition, across all levels of trait variation, and across all six different models. In this specific hierarchical trait-based HOI model, it was possible for a third species to alleviate the competitive dominance of a stronger competitor over a weaker competitor thereby facilitating species coexistence. For instance, a stronger competitor i.e., a species with lower trait value will easily dominate another species that has a higher trait value in accordance with the pairwise hierarchical function. In such a scenario, it was possible for the dominant species to outcompete a weaker competitor in the absence of a higher-order interaction. However, a presence of a third species and trait-based higher order interaction could significantly alleviate this dominance and thereby alleviate intense pairwise competition. This was possible when the third species had similar trait value as the dominant competitor, which then negates the stronger competitor dominance over the weaker competitor. Consequently, this could lead to species coexistence. As an example with three species (Fig. A2), we show exactly this.

However, at all levels of width in species competition, and different types of species interactions (whether pairwise or HOIs), higher individual variation does not promote species coexistence. Previous studies have shown the impact of HOIs from an ecological perspective where HOIs were drawn from random distributions (Bairey *et al*. 2016; Singh & Baruah 2021; Kleinhesselink *et al*. 2022), and the evolutionary context was largely ignored. In one of the models where we included evolutionary-HOIs, where phenotypic traits of species interacted through a Gaussian interaction kernel, stable coexistence of species was further impaired. In this specific model, higher-order effects were determined by species mean position in the one-dimensional trait axis. Pairwise trait-based species competition was impacted by the trait of a third species. If two species had similar mean trait values, pairwise species competition given by the Gaussian interaction kernel would be strong. The trait-based higher-order effects of the third species would be even stronger if the third species had similar position or similar value on the one-dimensional trait axis. At evolutionary equilibrium, this further led to low number of species coexisting. HOIs have previously been suggested to stabilize large number of species coexisting together. When evolutionary HOIs mediated through trait-based interactions come into play, species coexistence was disrupted irrespective of the presence of high or low individual variation. These evolutionary HOIs are low-dimensional and structured by their position in the trait axis. Consequently, in addition to pairwise trait-based species competition being low-dimensional, trait-based HOIs compound and structure species competition tightly in the one-dimensional trait axis further disrupting species coexistence. Although, empirical quantification of HOIs in natural ecosystems is difficult, ignoring such interactions would limit our understanding of the mechanisms behind species coexistence in complex communities. In comparison, our results indicate that low-dimensional trait-based HOI could promote species coexistence provided such HOI act in a hierarchical manner.

While modelling HOIs that were not dictated by mean phenotypic traits, but structured by random numbers, species coexistence can be stabilized but when species exhibited low individual trait variation. This happened when the strength of intraspecific HOIs was larger than interspecific HOIs. Such a formulation meant that HOIs were not mediated by any measurable aspect of phenotypic or functional traits but rather than by some other form. For instance, such HOIs could be assumed to be mediated through density or secondary traits. By encapsulating such a form through random numbers, we wanted to understand the conditions under which individual variation and HOIs together could impact the patterning of species coexistence. Our results demonstrated that HOIs that are higher in dimension, in contrast to only pairwise interactions or evolutionary HOIs, could lead to not only higher number of species coexisting together but also higher amount of trait patterning for a given amount of trait variation. In models of mixed HOIs, i.e., HOIs that had both positive or negative effects on pairwise species competition, coexistence was similarly disrupted when species had high trait variation similar to results from models of pairwise competition. Mixed HOIs were randomly sampled from a uniform distribution of *U*[-0.01, 0.01]. Their impact on species coexistence were negligible in contrast to other interaction models possibly due to two reasons. One reason could that effects of some positive HOIs are equally negated by negative HOIs such that their impact on trait patterning and coexistence was particularly dictated by pairwise species competition. Another reason could be due to the fact that higher-order effects of such HOIs were less strong in comparison to pairwise effects. When one increases the bounds of the uniform distribution from which such mixed HOIs were sampled, it realistically leads to uninformative abundances due to the HOI effects being linear w.r.t to the competitor species. A handful of positive HOIs from the mixed HOI distribution could lead to unrealistic species abundances, which puts a bound to how strong of positive HOIs one could sample (Singh & Baruah 2021).

In pure pairwise competitive communities, species coexistence will be disrupted as strength of competition increases, given a particular level of intraspecific variation. Similarly, as individual variation increases, given a particular level of competition, species coexistence will again be disrupted. Therefore, we should expect that any increase in individual variation would be counteracted by selection that decreases competitive interactions. This should lead to trait divergence over evolutionary time scales. Since, in our simulations we begin with saturated communities, as heritable individual variation increases, species will tend to evolve away from each other. In doing so they will encounter other species at other points in the trait axis more often than when individual variation is low. Consequently, coexistence between species will decrease in the case of high individual variation, and we report such a result, when the interactions are purely pairwise in nature as well as when trait-based HOIs are formulated by a Gaussian kernel.

Trait variation within species in a community is widely observed, but the implication of such trait variation on patterning of traits is still debated (Götzenberger *et al*. 2012). In our eco-evolutionary model, where competition between species included both pairwise and trait-mediated HOIs, increases in trait variation led to low trait clustering and more even spacing (Fig. 4). Lotka-Volterra models dominated by only pairwise interactions generally support the idea that species tend to distribute more evenly along a trait axis than expected by neutral evolution for the given trait (Barabás et al. 2012; Barabás and D’Andrea 2016). When there is high heritable variation in the trait, evolution would be faster compared to when there is low heritable variation. Thus, expectedly due to faster evolutionary dynamics caused by high heritable variation, species would move away from each other leading to lesser trait overlap and greater trait divergence in comparison to when heritable variation is low. When evolutionary HOIs came into play, trait divergence due to strong pairwise competition was further exacerbated particularly when all the three species involved in HOIs had similar mean phenotypic traits. This was understandable as the third species having similar mean trait value further negatively impacted species growth which led to higher divergence of species traits with lower number of species coexisting together. Evolutionary HOIs, thus, further promoted the ‘limiting similarity principle’ irrespective of the presence of high or low individual variation. Higher trait patterning, however, was seen only when intraspecific HOIs were modelled to be stronger than interspecific HOIs and when trait variation was low. As number of species that coexisted at evolutionary equilibrium increased, species generally got tightly packed along the one-dimensional trait axis. Naturally, if trait variation was high, the overlap of species traits increased, which led to fierce competition that could potentially lead to competitive exclusion. Competitive exclusion will also depend on other factors like width of the competition kernel. If the width of the competition kernel is high, say 0.3, species with dissimilar traits that overlap could still compete fiercely. Even in such scenarios, competitive exclusion could be avoided, and trait clustering could still be maintained provided a specific structure of high-dimensional HOIs are at play i.e., intraspecific HOIs should be stronger than interspecific HOIs. In other words, due to stronger intraspecific HOIs, species limited themselves more than they limited others which resulted in many coexisting species tolerating higher trait overlap. (Fig. 3, intraHOI, *ω =* 0.3). This could explain why many species are similar in functional trait space and could still coexist. The coexistence of a large number of competing species could be understood from the point that pairwise interactions and HOIs could be mediated through mechanisms that impact trait and population dynamics differently. In contrast, trait clustering in hierarchical models were slightly higher than all the other models compared. Despite that high trait variation still led low species clustering in the trait axis. Only at low competition levels i.e., *ω =* 0.1, trait clustering could be tolerated at high levels of trait variation.

Recent advances in understanding community stability when eco-evolutionary dynamics are at play, have shown that multispecies communities may be stable or unstable depending on the relative speed of evolutionary and ecological processes (Patel *et al*. 2018). That is, direct ecological or evolutionary, or indirect eco-evolutionary feedbacks will affect community stability depending on the relative timescales of ecological and evolutionary processes (Patel *et al*. 2018). Our results suggest that higher intraspecific variation could lead to robust species coexistence in the presence of HOIs (Fig. 4). However, this result could be due to the fact that higher trait variation leads to lower number of species coexisting which consequently could improve community robustness. In contrast, lower trait variation, particularly in models where intraspecific HOIs were stronger than interspecific HOIs (intraHOI), led to higher number of species coexisting with a significantly higher trait patterning. In such cases, robustness of species coexistence was much lower. High intraspecific variation led to faster evolutionary dynamics and the few species that coexisted separated themselves in a unidimensional trait axis both in the presence and absence of HOIs. Consequently, traits of persisting species had positions in the trait axis that could be advantageous for community robustness.

Our modelling results have the potential to indicate the possibility of HOIs in maintaining species coexistence despite differences in competitive ability. Evolutionary trait-mediated HOIs as modelled here can be one of the many mechanisms that could lead to distinct patterns of species coexistence. HOI terms could also be positive, indicating the possible facilitative effects of HOIs on species pairwise interactions. However, further development is needed to understand the effect of positive HOIs on species coexistence (Singh & Baruah 2021).

## Author contributions

GB formulated the study, analyzed the model, ran the simulations with help and contributions from GyB. GB, RJ, and GyB contributed to writing the manuscript. The authors declare no conflict of interest.

