## APPENDIX for "Evolutionary effects of individual variation and dimensionality of higher-order interactions on robustness of species coexistence"

Gaurav Baruah[a], Gyorgy Barabas[b], Robert John[c]

[a]*Faculty of Biology, Theoretical Biology, University of Bielefeld, 33501 Bielefeld, Germany.*
[b]*Department of Physics, Chemistry and Biology (IFM), Linkoping University, SE 58183 Linkoping, Sweden*
[c]*Department of Biological Sciences, Center for Climate and Environmental Studies, IISER Kolkata, India.*

---

*Keywords:* Stability, coexistence, robustness, trait clustering, intraspecific variation

---

### 1. Lotka-Volterra pairwise interaction model

We model the eco-evolutionary dynamics of S species that competes for resources in a community. Competition and growth of an individual of a species is dictated by a quantitative trait $z$ [4] . An individual of a species $i$ can be described by its quantitative trait value $z$ in a uni-dimensional trait axis. Accordingly, the distribution of the quantitative trait $z$ is Gaussian, $p(z)$, with mean $\mu_i$ for species $i$ and variance $\sigma_i$. If $N_i(t)$ is the number of individuals of species $i$ in time t, then $N_i(t)p_i(z,t)dz$ is the population density of species $i'$s individuals with trait value between $z$ and $z+dz$ [4].

Following Barabas et al, 2016, the per capita Lotka-Volterra growth rate for $S$ species in a competitive community dominated by pairwise interactions can be written as:

$$r(\vec{N},\vec{p},z) = b(z) - \sum_{j=1}^{S} N_j \int \alpha(z,z')p_j(z')dz', \tag{1}$$

where $b(z)$ is the instrinsic growth rate of trait $z$ and $\alpha(z,z')$ is the competition function that captures competition between trait $z$ and $z'$. The per captita growth rate in equation (1) is directly dependent on trait $z$. Competition among individuals of different species in the community also depends solely on the trait $z$. Further, the summation term in equation (1) ensures that every species in the community has an effect on the growth of species $i$ in the community. Our model is in the quantitative genetic limit [3] in the sense that the trait distribution $p(z)$ remains normal and only the mean of the distribution $\mu_i$ is subjected to selection due to competition and growth. Thus the variance and shape of $p(z)$ remains undisturbed.

### 2. Lotka-Volterra higher order interactions

With this pairwise Lotka-Volterra quantitative model, we introduce higher order interactions in equation (1) in the following way:

$$r(\vec{N},\vec{p},z) = b(z) - \sum_{j=1}^{S} N_j \int \alpha(z,z')p_j(z',t)dz' - \sum_{k=1}^{S}\sum_{j=1}^{S} N_k N_j \int\int \epsilon_k(z,z',z'')p_k(z',t)dz''p_j(z',t)dz', \tag{2}$$

Where $\epsilon_k(z,z')$ is the pair-wise interaction term of individuals of two different traits $z$ and $z'$ (in the same way as $\alpha(z,z')$) . However,the pairwise interaction between two individuals is modulated by the density of a third species $k$. The double summation ensures that each species modulates the pairwise interaction between two other species.

$\alpha(z,z')$ is the Gaussian competition kernel in (1) such that individuals with very similar trait will compete strongly whereas individuals that are far apart in trait values $z$ will compete weakly, given as:

$$\alpha(z,z') = \exp\left(\frac{-(z-z')^2}{\omega^2}\right),$$

Further,

$$\epsilon_k(z,z',z'') = \exp\left(\frac{-(z-z'-z')^2}{\omega^2}\right)$$

where $\epsilon_k(z,z,z'')$ captures the trait-mediated HOIs. Here, this particular formulation could also capture intraspecific HOIs where individuals belonging to the same species can modulate pairwise interactions between individuals belonging to the same species.

Growth rate of an individual with trait value $z$ is given by $b(z)$ which is a rectangular function along the trait axis given as:

$$b(z) = \begin{cases} 1, & \text{if } \theta \geq z \geq -\theta \\ 0, & \text{otherwise,} \end{cases}$$

Where $\theta$ is the limit of the trait axis such that any individual which has a trait value outside of the range of $[-\theta, \theta]$ will have zero growth.

With this growth equation, the dynamics of species $i$ can be written as :

$$\frac{dN_i(t)}{dt} = \int r_i(\vec{N}, z, t) p_i(z, t) dz, \tag{3}$$

Expanding equation (3) gives:

$$\frac{dN_i(t)}{dt} = N_i(t) \int \Big( b(z) - \sum_{j=1}^{S} N_j \int \alpha(z, z') p_j(z') dz' - \sum_{k=1}^{S} \sum_{j=1}^{S} N_k N_j \int \int \int \epsilon_k(z, z', z'') p_k(z', t) dz'' p_j(z', t) dz' \Big) p_i(z, t) dz,$$

$$\frac{dN_i(t)}{dt} = N_i(t) \Big( \int \big( b(z) p_i(z, t) dz \big) - \sum_{j=1}^{S} N_j \int \int \alpha(z, z') p_j(z', t) p_i(z, t) dz' dz - \sum_{k=1}^{S} \sum_{j=1}^{S} N_k N_j \int \int \epsilon_k(z, z', z'') p_k(z', t) dz' p_j(z', t) p_i(z, t) dz dz'' \Big),$$

Finally simple algebra of the above equations would lead to

$$\frac{dN_i(t)}{dt} = N_i(t) \Big( b_i(t) - \sum_{j=1}^{S} \alpha_{ij}(t) N_j(t) - \sum_{k=1}^{S} \sum_{j=1}^{S} \epsilon_{ijk}(t) N_j(t) N_k(t) \Big), \tag{4}$$

Where,

$$\alpha_{ij}(t) = \int \int \alpha(z, z') p_j(z', t) p_i(z, t) dz' dz = \frac{\omega}{\sqrt{2\sigma_i^2 + 2\sigma_j^2 + \omega^2}} \exp\big( \frac{-(\mu_i(t) - \mu_j(t))^2}{2\sigma_i^2 + 2\sigma_j^2 + \omega^2} \big). \tag{5}$$

The double integral is the weighted sum of interactions between individual of species $i$ which has $p_i(z)$ as its trait distribution and species $j$ which has $p_j(z')$ as its trait distribution. This term quantifies the competition species $i$ faces from species $j$. And,

$$b_i(t) = \int b(z) p_i(z, t) dz = \frac{1}{2} \Bigg[ erf(\frac{\theta - \mu_i}{\sqrt{2}\sigma_i}) + erf(\frac{\theta + \mu_i}{\sqrt{2}\sigma_i}) \Bigg]. \tag{6}$$

$$\epsilon_{ijk}(t) = \int \int \int \epsilon_k(z, z', z'') p_j(z', t) p_i(z, t) p_k(z'', t) dz dz' dz'' = \frac{\omega}{\sqrt{2\sigma_i^2 + 2\sigma_j^2 + 2\sigma_k^2 + \omega^2}} \exp\big( \frac{-(\mu_i(t) - \mu_j(t) - \mu_k(t))^2}{2\sigma_i^2 + 2\sigma_j^2 + 2\sigma_k^2 + \omega^2} \big) \tag{7}$$

The above term captures the three-way strength of interaction. Pairwise competition between species $i$ with mean trait $\mu_i$ and species $j$ with mean trait $\mu_j$ is influenced by the density of another species $k$ [7]. Finally the dynamics of the trait of a species in response to growth as well as in response to interspecific competition due to pairwise interactions among species and higher-order interactions can be formulated as ,

$$\frac{d\mu_i(t)}{dt} = h_i^2 \int (z - \mu_i(t)) r_i(\vec{N}, z, t) p_i(z, t) dz, \tag{8}$$

where, $h_i^2$ is the fraction of heritable variation of the phenotype $\mu_i$ of species $i$. In our simulations we fixed the heritable variation to 0.1 for all the species. This can be further analytically integrated and expanded as :

$$\frac{d\mu_i(t)}{dt} = h_i^2 \sigma_i^2 \Big( \bar{g_i}(t) - \sum_{j=1}^{S} \beta_{ij}(t) N_j(t) - \sum_{k=1}^{S} \sum_{j=1}^{S} \gamma_{ijk}(t) N_j(t) N_k(t) \Big), \tag{9}$$

where first term of the above equationis:

$$\bar{g_i}(t) = \int \big((z-\mu_i(t))b(z)p_i(z,t)dz\big) = \frac{1}{\sqrt{2\pi}\sigma_i}\Bigg[\exp\big(\frac{-(\theta+\mu_i)^2}{2\sigma_i^2}\big) - \exp\big(\frac{-(\theta-\mu_i)^2}{2\sigma_i^2}\big)\Bigg],$$

Similarly,

$$\beta_{ij}(t) = \int\int(z-\mu_i(t))\alpha(z,z')p_j(z',t)p_i(z,t)dz'dz = \frac{2\omega(\mu_j(t)-\mu_i(t))}{\sqrt[3]{2\sigma_i^2+2\sigma_j^2+\omega^2}}\exp\big(-\frac{(\mu_i(t)-\mu_j(t))^2}{2\sigma_i^2+2\sigma_j^2+\omega^2}\big)$$

And,

$$\gamma_{ijk} = \int\int\int(z-\mu_i(t))\epsilon_k(z,z',z'')p_j(z',t)p_i(z,t)p_k(z'',t)dzdz'dz'' =$$
$$\frac{2\omega(\mu_j(t)-\mu_i(t)-\mu_k(t))}{\sqrt[3]{2\sigma_i^2+2\sigma_j^2+2\sigma_k^2+\omega^2}}\exp\big(-\frac{(\mu_i(t)-\mu_j(t))^2}{2\sigma_i^2+2\sigma_j^2+2\sigma_k^2+\omega^2}\big).$$

Thus ,

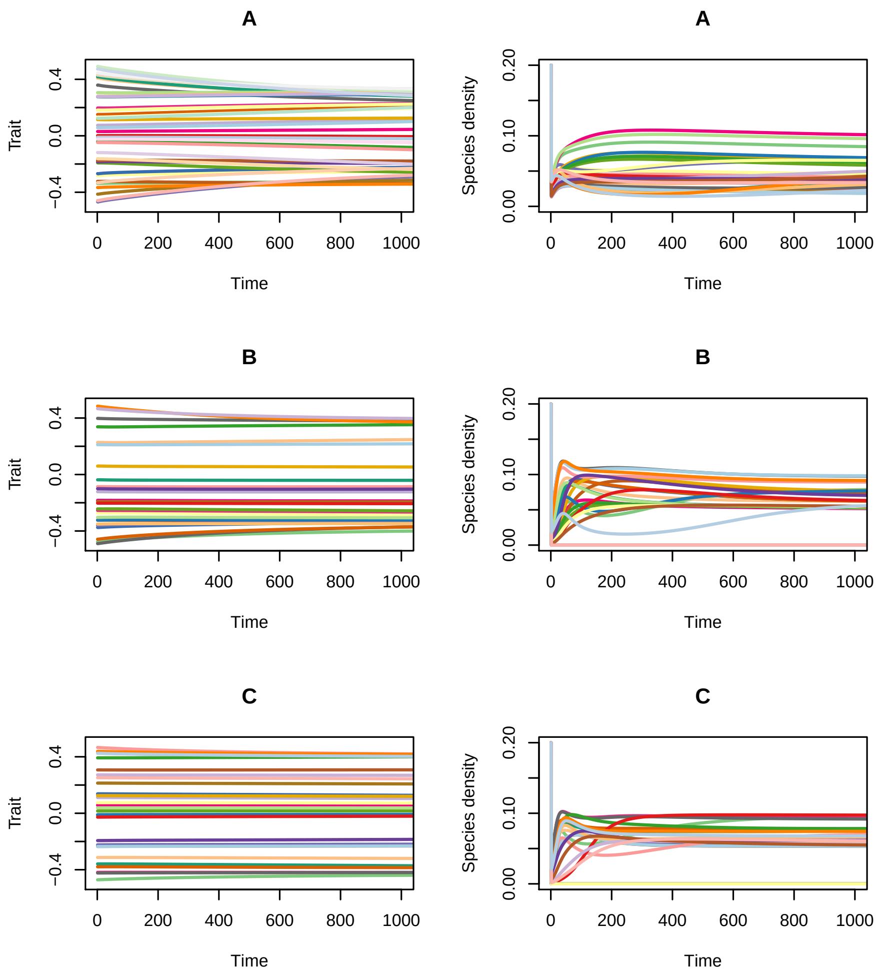


Fig.A 1: Example dynamics of trait (left column) and species density (right column) over time for three levels of intraspecific variation for a particular w = 0.3. (A) High intraspecific variation; (B) Medium intraspecific variation; (c) low intraspecific variation. Different colors represent different species.

$$\frac{d\mu_i(t)}{dt} = h_i^2\sigma_i^2\Big(\bar{g_i}(t) - \sum_{j=1}^{S}\beta_{ij}(t)N_j(t) - \sum_{k=1}^{S}\sum_{j=1}^{S}\gamma_{ijk}(t)N_j(t)N_k(t)\Big). \quad (10)$$

From equation 9, the first term denotes evolutionary pressure due to growth of the trait in the trait axis, the second term $\beta_{ij}$ denotes the selection acting on the mean trait value of species $i$ due to pairwise competition with other species in the trait axis, and the third term $\gamma_{ijk}$ indicates the selection that acts on the trait due to higher-order interactions in the trait axis.

### 3. Hierarchical evolutionary higher-order equations

We model another trait-mediated HOI where interactions among species are mediated through mean phenotypic traits. However, the per-capita growth rate of a species follows a different functional form. Intrinsically, species trait value in the one-dimensional trait axis define how competitively dominant they are which then trades off with their ability to grow. In this particular model, lower trait values correspond to stronger competitors but with high mortality and higher trait values correspond to weaker competitors with low mortality. This per-capita growth rate of a phenotype $z$ belonging to species $i$ can be written as:

$$r(\vec{N},\vec{p},z) = b(z) - \sum_{j=1}^{S} N_j \int \alpha(z,z')p_j(z',t)dz' - \sum_{k=1}^{S}\sum_{j=1}^{S} N_k N_j \int\int \epsilon(z,z',z'')p_k(z',t)dz''p_j(z',t)dz', \tag{11}$$

Here,

$$b(z) = 1 - \exp(-z-\theta), \tag{12}$$

$$\alpha(z,z') = \frac{1}{2}\Big(\tanh(\frac{(z-z')}{w})+1\Big), \tag{13}$$

And,

$$\epsilon(z,z',z'') = \frac{1}{2}\Big(\tanh(\frac{(z-(z'+z''-z_0))}{\omega_2})+1\Big) \tag{14}$$

Population and evolutionary trait dynamics can easily be calculated as equation 3 and equation 8. Note that the integrals are not analytically solvable and we use simpler numerical methods to approximate the integrals. Equation 12 that growth rate falls when as trait value decreases. However, individuals are stronger competitors (equation 13). IN all our simulations $z_0 = 0.7$, $\theta = 0.5$ and $\omega_2 = 0.1$.

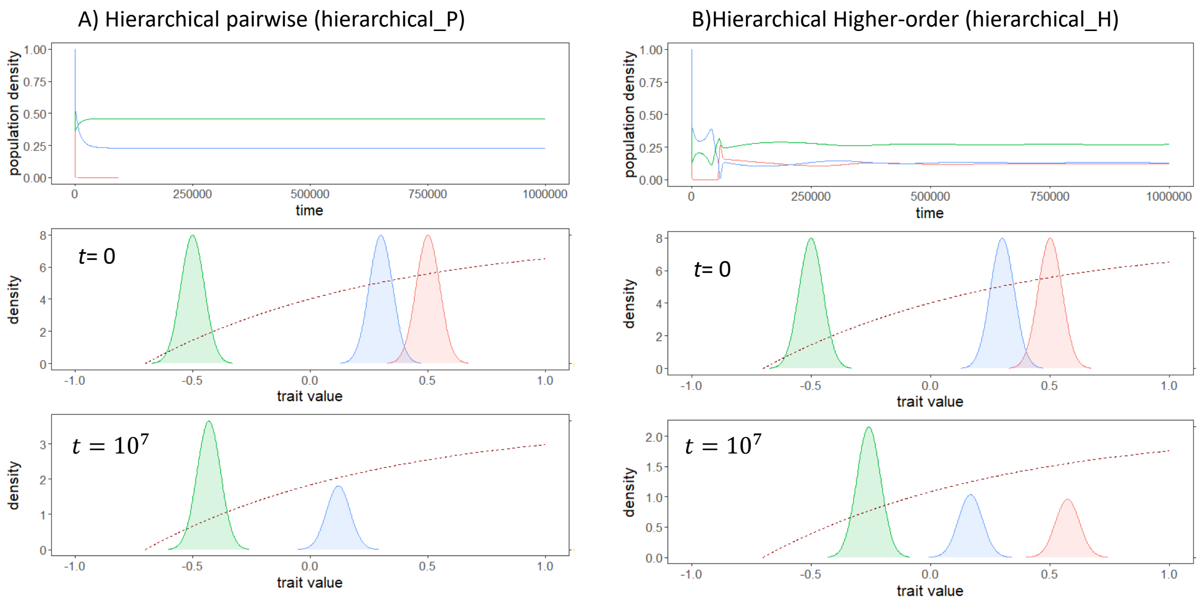


Fig.A 2: Dynamics of three species engaged in pairwise hierarchical competition (A) and higher-order interaction (B). A) Species with lower trait values are more dominant in terms of competition than species with higher trait values. This comes with a trade-off in growth rate. At the start of the simulation, three species are placed in the trait-axis with same trait variance and with dominance hierarchy.Naturally at $t = 10^7$ the weakest competitor goes extinct. B) In the presence of trait-based HOI, this scenario changes completely and all three species coexist. This was because the impact of the most dominant competitor on the weakest competitor was alleviated by the third species resulting in coexistence of all three species. Initial abundance of all species was fixed to 1 and trait variance of all species was fixed at 0.05. $z_0$ was fixed at 0.7, and strength of pairwise hierarchical interaction $\omega = 0.3$ and strength of HOI $\omega_2 = 0.1$.

### 4. Jacobian, stability and robustness of species coexistence

The Jacobian of a dynamical system at a given point is given as:

$$J_{ij} = \frac{\partial\big(\frac{dN_i(t)}{dt}\big)}{\partial N_j}$$

.

At eco-evolutionary equilibrium, when changes in the abundances of species is zero which is the equilibrium condition of S species coexisting, then we can write from equation (3) that :

$$\frac{dN_i(t)}{dt} = \Big(b_i - \sum_{j=1}^{S} \alpha_{ij} N_j(t) - \sum_{k=1}^{S}\sum_{j=1}^{S} \epsilon_{ijk} N_j(t) N_k\Big) = 0, \tag{15}$$

which means at equilibrium, species $i$ will follow the equation below :

$$\Big(b_i = \sum_{j}^{S} \alpha_{ij} N_j - \sum_{k}^{S}\sum_{j}^{S} \epsilon_{ijk} N_j N_k\Big). \tag{16}$$

Evaluating the Jacobian at any time point t:

$$J_{ij} = \frac{\partial\big(\frac{dN_i(t)}{dt}\big)}{\partial N_j} = \delta_{ij}\Big(b_i - \sum_{j}^{S} \alpha_{ij} N_j - \sum_{k}^{S}\sum_{j}^{S} \epsilon_{ijk} N_j N_k\Big) - N_i \alpha_{ij} - N_i\Big(\sum_{k}^{S} \epsilon_{ijk} + \sum_{k}^{S} \epsilon_{ikj}\Big) \tag{17}$$

where $\delta_{ij}$ is the Kronecker delta where

$$\delta_{ij} = \begin{cases} 1, & \text{if } i = j \\ 0, & \text{otherwise.} \end{cases}$$

Since we are assuming that at the end of our simulations communities have reached equilibrium, the kronecker delta becomes zero. Hence equation 17 can be written as

$$J_{ij} = \frac{\partial\big(\frac{dN_i(t)}{dt}\big)}{\partial N_j} = -N_i \alpha_{ij} - N_i\Big(\sum_{k}^{S} \epsilon_{ijk} + \sum_{k}^{S} \epsilon_{ikj}\Big) \tag{18}$$

Equation (18) is a modified community matrix that incorporates higher-order 3 way interaction [6]. From this equation at particular equilibrium time point, one can estimate eigenvalues from this Jacobian matrix and in turn can estimate stability and average robustenss of species coexistence at equilibrium.

If the eigenvalues of the equation (18) yields all negative eigenvalues, then the community at that point is stable. In other words, suppose, S species coexist at the end of a simulation run, and if the eigenvalues of the Jacobian at that time point has all S negative eigenvalues then the equilibrium is locally stable.

Thus local stability of an ecological community will be guaranteed if the Jacobian matrix in equation (18) has eigenvalues that have negative real parts. This means that any perturbation at that point will decay along its eigendirection with a rate equal to the eigenvalue. The robustness of this local stability of an ecological community is given by the determinant of the Jacobian matrix [1, 5, 2]. As explicitly formulated in Barabas et al 2015, it is given by geometric mean of the absolute values of the eigenvalues as :

$$\sqrt[S]{|\lambda_1||\lambda_2|...|\lambda_S|} = \Big(\prod_{i=1}^{S} |\lambda_i|\Big)^{\frac{1}{S}} = \exp(\frac{1}{S}\log(\prod_{i=1}^{S} |\lambda_i|) = \exp(\bar{\log}(|\lambda|))$$

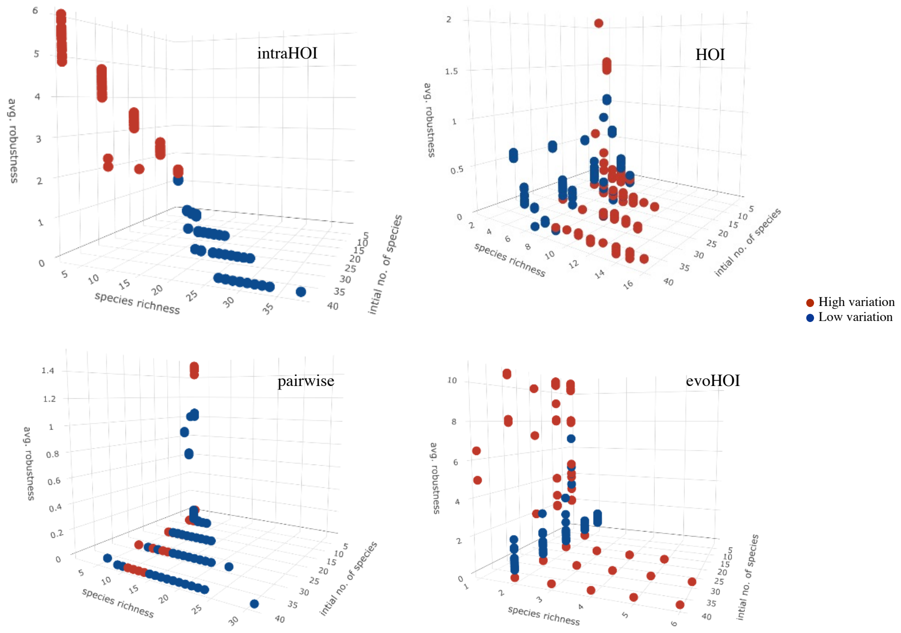


Fig.A 3: Average robustness of species coexistence plotted against species richness, and initial number of species in the community for four different interaction models namely intraHOI, HOI, pairwise, evoHOI. Generally, individual trait variation leads to high robustness of species coexistence only when species coexistence was low.

[5] Levins, R. 1979. Coexistence in a Variable Environment. The American Naturalist 114:765–783. ISSN 00030147, 15375323. URL `http://www.jstor.org/stable/2460550`.

[6] May, R. M. 1973. Qualitative Stability in Model Ecosystems. Ecology 54:638–641. URL `http://doi.wiley.com/10.2307/1935352`.

[7] Terry, J. C. D., Morris, R. J., and Bonsall, M. B. 2018. Trophic interaction modifications disrupt the structure and stability of food webs. bioRxiv page 345280. URL `https://www.biorxiv.org/content/10.1101/345280v2`.
